# Modulation of dorsal premotor cortex disrupts neuroplasticity of primary motor cortex in young and older adults

**DOI:** 10.1101/2023.05.28.542670

**Authors:** Wei-Yeh Liao, George M. Opie, Ulf Ziemann, John G. Semmler

**Author notes:** Correspondence: Wei-Yeh Liao School of Biomedicine The University of Adelaide Adelaide, South Australia 5005 Australia Telephone: Int.

## Abstract

Although transcranial magnetic stimulation (TMS) research demonstrates that dorsal premotor cortex (PMd) influences neuroplasticity within primary motor cortex (M1), it is unclear how ageing modifies this communication. The present study investigated the influence of PMd on different indirect (I) wave inputs within M1 that mediate cortical plasticity in young and older adults. 15 young and 15 older participants completed two experimental sessions that examined the effects of intermittent theta burst stimulation (iTBS) to M1 when preceded by iTBS (PMd iTBS-M1 iTBS) or sham stimulation (PMd sham-M1 iTBS) to PMd. Changes in corticospinal excitability post-intervention were assessed with motor evoked potentials (MEP) recorded from right first dorsal interosseous using posterior-anterior (PA) and anterior-posterior (AP) current single-pulse TMS (PA*_1mV_*; AP*_1mV_*; PA*_0.5mV_*, early I-wave; AP*_0.5mV_*, late I-wave). Although PA*_1mV_* did not change post-intervention (*P* = 0.628), PMd iTBS-M1 iTBS disrupted the expected facilitation of AP*_1mV_* (to M1 iTBS) in young and older adults (*P* = 0.002). Similarly, PMd iTBS-M1 iTBS disrupted PA*_0.5mV_* facilitation in young and older adults (*P* = 0.030), whereas AP*_0.5mV_* facilitation was not affected in either group (*P* = 0.218). This suggests that while PMd specifically influences the plasticity of early I-wave circuits, this communication is preserved in older adults.

## Introduction

One of the universal effects of ageing is widespread deficits in motor function. Although these deficits occur at all levels of the motor system, the structural, functional, and biochemical changes within the brain are important (Seidler *et al*., 2010). In particular, alterations to the ability of the brain’s motor system to continuously modify its structure and function are a critical factor. Termed neuroplasticity, this process is initially mediated by changes in the strength of synaptic communication with long-term potentiation (LTP) and depression (LTD), and underpins the ability to learn new motor skills (Buonomano & Merzenich, 1998; Sanes & Donoghue, 2000). While the capacity for neuroplastic change is present across the lifespan, some studies using non-invasive brain stimulation (NIBS) show reduced plasticity in older adults (Müller-Dahlhaus *et al*., 2008; Fathi *et al*., 2010; Todd *et al*., 2010; Freitas *et al*., 2011). This reduced plasticity may contribute to the motor deficits that limit the ability of older adults to learn new motor skills that may be essential for daily life. However, the neurophysiological mechanisms underpinning these changes with advancing age remain unclear.

Transcranial magnetic stimulation (TMS) is a type of NIBS that allows investigation of specific neuronal networks within the motor system with high temporal resolution. Application of TMS over primary motor cortex (M1) produces a complex series of descending volleys within corticospinal neurons that summate at the spinal cord, resulting in a motor evoked potential (MEP) in targeted muscles (Di Lazzaro *et al*., 1998; Rossini *et al*., 2015). The first of these waves likely represents direct activation of corticospinal neurons, whereas subsequent waves are thought to reflect the indirect activation of interneuronal inputs to the corticospinal neurons (Di Lazzaro *et al*., 2012; Ziemann, 2020). These responses are referred to as indirect (I) waves and are named early (I*_1_*) or late (I*_2_,* I*_3_*) based on the order of their appearance, which occurs with a periodicity of ∼1.5 ms (Di Lazzaro *et al*., 2012; Ziemann, 2020). Early and late I-waves can be preferentially recruited by applying low-intensity single-pulse TMS with different current directions (Sakai *et al*., 1997; Di Lazzaro *et al*., 2001; Ni *et al*., 2010). For example, a posterior-anterior (PA) current (relative to the central sulcus) preferentially recruits early I-waves, whereas an anterior-posterior (AP) current preferentially recruits late I-waves (Sakai *et al*., 1997; Di Lazzaro *et al*., 2001; Ni *et al*., 2010). Using these measures, previous work has shown that the ability to recruit late I-waves predicts the response to plasticity-inducing TMS paradigms over M1 (Hamada *et al*., 2013; Wiethoff *et al*., 2014) and that the late I-waves are behaviourally relevant to the acquisition of fine motor skills (Hamada *et al*., 2014).

I-wave circuits are also involved in mediating the communication between other motor nodes and M1 (Groppa *et al*., 2012; Volz *et al*., 2015; Spampinato *et al*., 2020; Opie *et al*., 2022; Casarotto *et al*., 2023), which form a wider network that influences M1 plasticity and learning (Huang *et al*., 2018; Liao *et al*., 2022). In particular, the dorsal premotor cortex (PMd) facilitates the planning, prediction, and correction of movements during motor learning by updating the activity of M1 (Chouinard *et al*., 2005; Nowak *et al*., 2009; Parikh & Santello, 2017). Previous studies have demonstrated that the application of repetitive TMS (rTMS) techniques (such as theta burst stimulation; TBS) over PMd is able to modify M1 excitability, plasticity, and motor skill acquisition (Huang *et al*., 2018; Meng *et al*., 2020).

Furthermore, while PMd influences both early and late I-wave excitability (Liao *et al*., 2023), there is a stronger effect on the late I-waves (Volz *et al*., 2015; Aberra *et al*., 2020). Taken together, it is likely that the influence of late I-waves on M1 plasticity reflects inputs from PMd.

Given the role of late I-wave circuits in mediating PMd-M1 communication, changes in late I-wave activity may affect the influence of PMd on M1 plasticity. In particular, late I-wave activity is known to be altered with advancing age (Opie *et al*., 2018). Age-related changes in I-wave excitability have been investigated using the paired-pulse TMS protocol short intracortical facilitation (SICF) (Opie *et al*., 2018), which revealed reduced I-wave excitability and a specific delay in the temporal characteristics of the late I-waves in older adults (Opie *et al*., 2018). Importantly, this delay influences NIBS-induced plasticity and is associated with specific aspects of motor behaviour in older adults (Opie *et al*., 2018; Opie *et al*., 2020). In addition, it is also known that PMd-M1 effective connectivity (Ni *et al*., 2015) and direct PMd modulation of early I-waves within M1 is reduced in older adults (Liao *et al*., 2023). Consequently, it is possible that the influence of PMd on M1 plasticity is altered with advancing age, but this remains to be tested.

The purpose of the present study was, therefore, to investigate the influence of PMd on the plasticity of early and late I-wave circuits in M1 of young and older adults. Given that previous work has used TBS to modulate M1 plasticity in young adults (Huang *et al*., 2018), we applied intermittent TBS (iTBS) over PMd in young and older participants and assessed how this influenced the neuroplastic response of M1 to iTBS. Different I-wave circuits were assessed by varying the direction of current used to apply TMS over M1. Although we expected iTBS over PMd to selectively modulate the plasticity of late I-wave circuits, we hypothesised that the effect of PMd on M1 plasticity would be weaker in older adults, given the likely alterations in late I-wave activity and PMd-M1 connectivity with advancing age.

## Materials and Methods

### Sample Size and Participants

15 young (mean ± standard deviation, 24.7 ± 5.0 years; range, 19-36 years) and 15 older adults (67.2 ± 5.4 years; 61-78 years) were recruited for the study via advertisements placed on notice boards within The University of Adelaide and the wider community, in addition to social media platforms. Applicants for the study were excluded if they had a history of psychiatric or neurological disease, current use of medication that affect the central nervous system, pregnancy, metal implants, or left handedness, as assessed by a standard TMS screening questionnaire (Rossi *et al*., 2011). The experiment was conducted in accordance with the Declaration of Helsinki and was approved by The University of Adelaide Human research Ethics Committee (H-026-2008). Subjects provided written, informed consent prior to participation.

### Experimental Arrangement

All participants attended two experimental sessions where iTBS or sham iTBS was applied to PMd, followed 30 minutes later by plasticity induction within M1 via iTBS (PMd iTBS-M1 iTBS, PMd sham-M1 iTBS). The same experimental protocol was used in both sessions (Fig. 1), with the order of intervention randomised between participants, and a washout period of at least 1 week was used between sessions. As diurnal variations in cortisol are known to influence the neuroplastic response to TMS (Sale *et al*., 2008), all sessions were completed between 11 am and 5 pm at approximately the same time of day for each participant.

**Figure 1.**
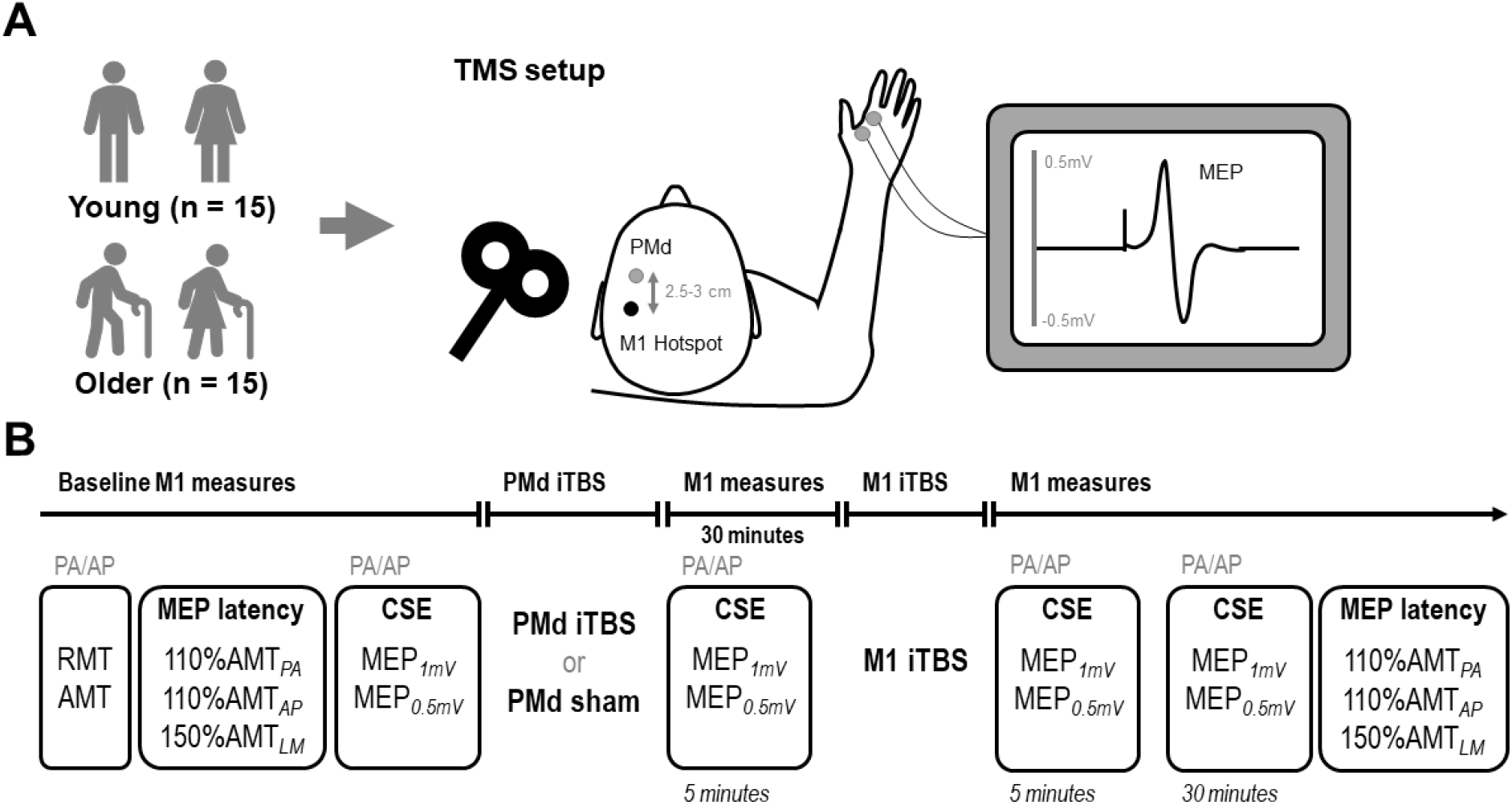
(A) Subject sample and experimental setup. (B) Experimental procedure. PA, posterior-to-anterior; AP, anterior-to-posterior; LM, lateral-to-medial; RMT, resting motor threshold; AMT, active motor threshold; MEP*_1mV_*, standard MEP of ∼ 1mV at baseline; MEP*_0.5mV_*, MEP of ∼ 0.5 mV at baseline; PMd, dorsal premotor cortex; iTBS, intermittent theta burst stimulation; CSE, corticospinal excitability.

During each experimental session, participants were seated in a comfortable chair with their hands resting and relaxed. Surface electromyography (EMG) was recorded from the first dorsal interosseous (FDI) of the right hand using two Ag-AgCl electrodes arranged in a belly-tendon montage on the skin above the muscle, with a third electrode attached above the styloid process of the right ulnar used to ground the electrodes. EMG signals were amplified (300x) and filtered (band-pass 20 Hz – 1 kHz) using a CED 1902 signal conditioner (Cambridge Electronic Design, Cambridge, UK) before being digitised at 2 kHz using a CED 1401 analogue-to-digital converter. Signal noise associated with mains power was removed using a Humbug mains noise eliminator (Quest Scientific, North Vancouver, Canada). EMG signals were stored on a PC for offline analysis. Real-time EMG signals were displayed on an oscilloscope placed in front of the participant to facilitate muscle relaxation during the experiment.

### Experimental Procedures

#### Transcranial magnetic stimulation (TMS)

A branding iron coil connected to two Magstim 200^2^ magnetic stimulators (Magstim, Whitland, UK) via a BiStim unit was used to apply TMS to left M1. The coil was held tangentially to the scalp at an angle of 45° to the sagittal plane, inducing a PA current relative to the central sulcus. The M1 hotspot was identified as the location producing the largest and most consistent MEPs within the relaxed FDI muscle of the right hand (Rossini *et al*., 2015). This location was marked on the scalp for reference and continuously monitored throughout each experimental session. All baseline, post-PMd iTBS, and post-M1 iTBS (5 minutes, 30 minutes) TMS was applied at a rate of 0.2 Hz, with a 10% jitter between trials to avoid anticipation of the stimulus.

Resting motor threshold (RMT) was recorded as the lowest stimulus intensity producing an MEP amplitude ≥ 50 µV in at least 5 out of 10 trials during relaxation of the right FDI. RMT was assessed at the beginning of each experimental session and expressed as a percentage of maximum stimulator output (% MSO) (Rossini *et al*., 2015). Active motor threshold (AMT) was then assessed, defined as the lowest % MSO producing an MEP amplitude ≥ 200 µV in at least 5 out of 10 trials during concurrent low-level activation (∼10% voluntary activation) of the right FDI (Hamada *et al*., 2013). These measures were then repeated using the AP current by rotating the coil 180°. Then, the stimulus intensities producing a standard MEP amplitude approximating 1 mV (MEP*_1mV_*; PA*_1mV_*, AP*_1mV_*), in addition to an MEP amplitude approximating 0.5 mV (MEP*_0.5mV_*; PA*_0.5mV_*, AP*_0.5mV_*), when averaged over 20 trials, were identified. The same intensities (MEP*_1mV_*, MEP*_0.5mV_*) were then applied following PMd iTBS and following M1 iTBS to assess changes in corticospinal excitability.

#### I-wave recruitment

To investigate the ability to recruit I-waves, the onset latencies of PA (early) and AP (late) MEPs were assessed relative to the MEP onset generated by direct activation of corticospinal neurons using a lateral-to-medial (LM) current (Hamada *et al*., 2013). A block of 15 MEP trials in the active FDI was recorded for 110% of AMT*_PA_* and AMT*_AP_*, in addition to 150% AMT*_LM_* (Hamada *et al*., 2013). If 150% AMT*_LM_* exceeded 100% MSO, 100% MSO was used, or if 150% AMT*_LM_* was below 50% MSO, 50% MSO was used (Hamada *et al*., 2013). The difference in mean onset latencies between PA and LM (PA-LM) and AP and LM (AP-LM) were calculated as measures of early and late I-wave recruitment efficiency, respectively (Hamada *et al*., 2013). In an attempt to reduce the confounding influence of muscle contraction on neuroplasticity induction (Huang *et al*., 2008; Thirugnanasambandam *et al*., 2011; Goldsworthy *et al*., 2015), these measures were recorded at the start and at the end of the experimental session, at least 45 minutes apart from the plasticity induction of PMd and M1.

#### Theta burst stimulation (TBS)

Intermittent theta burst stimulation (iTBS) was delivered over left PMd and left M1 using a Magstim Super-rapid stimulator (Magstim, Whitland, UK), connected to an air-cooled figure-of-eight coil. The coil was held tangentially to the scalp, at an angle of 45° to the sagittal plane, with the handle pointing backwards and laterally, inducing a biphasic pulse with an initial PA current followed by an AP return current (Suppa *et al*., 2008). In accordance with existing literature, iTBS consisted of bursts of three pulses given at a frequency of 50 Hz. Each burst was repeated at 5 Hz for 2 s, and repeated every 8 s for 20 cycles, totalling 600 pulses (Huang *et al*., 2005; Huang *et al*., 2008; Huang *et al*., 2018; Meng *et al*., 2020). The location of left PMd was defined as 8% of the distance between the nasion and inion (approximately 2.5 – 3 cm) anterior to the M1 hotspot, consistent with previous work (Münchau *et al*., 2002; Koch *et al*., 2007; Huang *et al*., 2018; Meng *et al*., 2020). The location of both the M1 hotspot and left PMd site were logged relative to the MNI-ICBM152 template using Brainsight neuronavigation (Rogue Research, Montreal, Quebec, Canada). These locations were then used to guide the assessment of RMT (RMT*_Rapid_*) over M1 with the Magstim Super-rapid stimulator, in addition to the application of iTBS over left PMd and M1 at 70% RMT*_Rapid_*.

Sham iTBS to left PMd was delivered using a sham figure-of-eight coil (replicating the coil click), with a bar electrode connected to a constant current stimulator (Digitimer, Hertfordshire, UK) placed underneath the coil delivering electrical stimulation (1.5 mA) to the scalp in order to mimic the pulse sensation. Following either intervention, participants provided answers to a visual analogue scale (VAS) questionnaire indexing the degree of discomfort, muscle activation, and localisation of scalp sensation during PMd iTBS.

### Data Analysis

Visual inspection of EMG data was completed offline, with any trials obtained from the resting muscle having EMG activity exceeding 25 µV in the 100 ms prior to stimulus application excluded from analysis (approximately 6.8% removed). The amplitude of MEPs obtained from resting muscle recordings was measured peak-to-peak and expressed in mV. The MEP onset latencies obtained from active muscle recordings was assessed with a semi-automated process using a custom script within the Signal program (v 6.02, Cambridge Electronic Design) and expressed in ms. MEP latency was recorded as the period from stimulus application to the resumption of voluntary EMG activity. This was defined as the point at which post-stimulus EMG amplitude exceeded the mean EMG amplitude recorded within the 100 ms pre-stimulus, plus 2 standard deviations. MEP onset latencies were averaged over individual trials within each subject and coil orientation. Within each participant, the mean LM MEP latencies were subtracted from the mean PA and AP MEP latencies to determine PA-LM and AP-LM MEP latency differences. Following TBS interventions, changes in MEP latency differences were quantified by expressing the post-intervention responses as a percentage of the baseline responses. Changes in MEP amplitude due to PMd iTBS were quantified by expressing post-PMd iTBS responses as a percentage of baseline MEP amplitude. For post-M1 iTBS, changes in MEP amplitude were quantified by expressing post-M1 iTBS responses as a percentage of post-PMd iTBS responses.

### Statistical Analysis

Visual inspection and Kolmogorov-Smirnov tests of the data residuals revealed non-normal, positively-skewed distributions for all TMS data. Consequently, generalised linear mixed models (GLMM’s), which can account for non-normal distributions (Lo & Andrews, 2015; Puri & Hinder, 2022), were used to perform all statistical analyses. Each model assessing MEP amplitude included single trial data with repeated measures and was fitted with Gamma distributions (Puri & Hinder, 2022), with all random subject effects included (intercepts and slopes) (Barr *et al*., 2013). Identity link functions were used for baseline MEP amplitude and latency differences while log link functions were used for post-iTBS normalised MEP amplitude and latency differences (Lo & Andrews, 2015; Puri & Hinder, 2022). To optimise model fit, we tested different covariance structures and the structure providing the best fit (assess with the Bayesian Schwartz Criterion; BIC) within a model that was able to converge was used in the final model. Two-factor GLMMs were used to compare effects of session (PMd iTBS-M1 iTBS, PMd sham-M1 iTBS) and age (young, older) at baseline in eight separate models for PA*_0.5mV_*, AP*_0.5mV_*, PA*_1mV_*, and AP*_1mV_* stimulation intensities and MEP amplitude. A three-factor model was used to compare the effects of session, age, and orientation (PA, AP) on PA-LM and AP-LM latency differences at baseline.

Changes in corticospinal excitability following PMd iTBS were investigated by assessing effects of session and age in four separate models for baseline-normalised PA*_1mV_*, AP*_1mV_*, PA*_0.5mV_*, and AP*_0.5mV_* MEP amplitude. Changes in corticospinal excitability following PMd iTBS-M1 iTBS and PMd sham-M1 iTBS were investigated by assessing effects of session, time (5 minutes, 30 minutes) and age in four separate models for PA*_1mV_*, AP*_1mV_*, PA*_0.5mV_*, and AP*_0.5mV_* MEP amplitude normalised to the mean post-PMd iTBS MEP amplitude. As AP*_1mV_*baseline stimulation intensities varied between sessions (see Table 1), AP*_1mV_* stimulation intensities were also included in the model as a covariate to assess if varying stimulation intensities confounded changes in post-intervention AP*_1mV_* MEP amplitude. Changes in I-wave recruitment following the intervention were investigated by assessing effects of session, age, and coil orientation on baseline-normalised average PA-LM and AP-LM latency differences. For all models, investigation of main effects and interactions were performed using custom contrasts with the Bonferroni correction, and significance was set at *P* < 0.05. Data for all models are presented as estimated marginal means (EMMs) and 95% confidence intervals (95% CI), whereas pairwise comparisons are presented as the estimated mean difference (EMD) and 95% CI for the estimate.

**Table 1.**
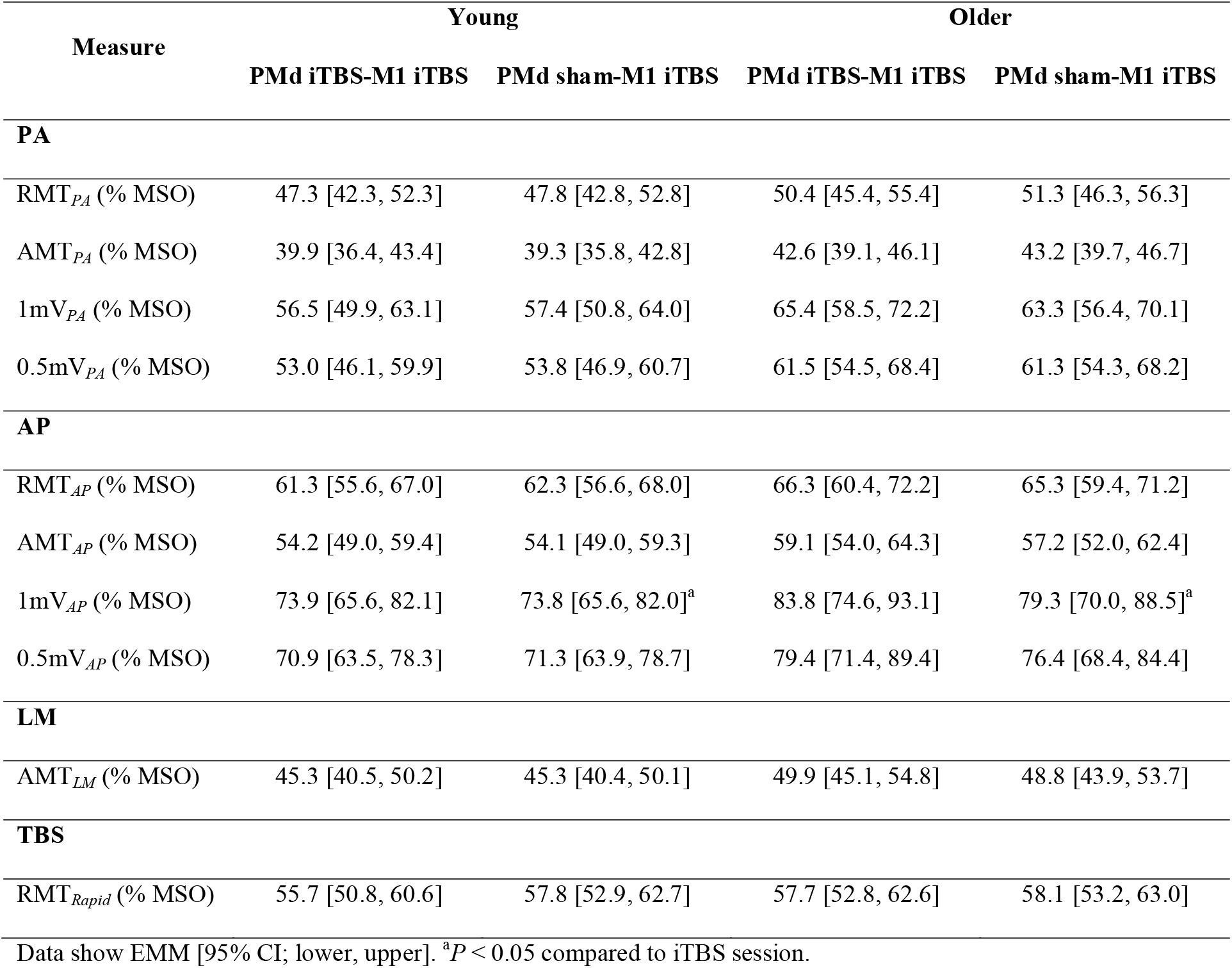
Baseline TMS intensities between sessions for young and older adults.

Furthermore, we used Spearman’s rank order correlation analysis to assess the relationship between different variables. Specifically, baseline MEP latency differences were correlated with changes in corticospinal excitability immediately following PMd iTBS to investigate if the ability to recruit I-waves is related to changes in corticospinal excitability. Baseline MEP latency differences were also correlated with changes in corticospinal excitability during the PMd sham-M1 iTBS session to investigate if the ability to recruit I-waves is related to changes in corticospinal excitability following M1 iTBS. In addition, changes in corticospinal excitability following PMd iTBS were also correlated with changes in corticospinal excitability following M1 iTBS (during PMd iTBS-M1 iTBS) to investigate if direct PMd modulation of M1 excitability is related to changes in M1 plasticity. Correlations are presented as Spearman’s ρ with false discovery rate-adjusted *P*-value of 0.05 following the Benjamini-Hochberg procedure. Lastly, differences in the perception of discomfort, extent of FDI activation, and localisation of stimulus during PMd iTBS and PMd sham were investigated by comparing VAS responses using paired t-tests with Bonferroni correction (*P* < 0.0167), with data presented as mean ± standard deviation.

## Results

All participants completed both experimental sessions without adverse reactions. We were unable to record PA*_1mV_* in one older male participant, AP*_0.5mV_* in two older participants (1 female, 1 male), and AP*_1mV_* in five participants (1 young female; 3 older females, 1 older male) due to high thresholds of activation (mean RMT*_PA_* = 80.0% MSO, mean RMT*_AP_* = 73.0% MSO). Baseline stimulation intensities are presented in Table 1. Stimulation intensities for AP*_1mV_* differed between sessions (*F*_1,46_ = 4.17, *P* = 0.047), with *post-hoc* comparisons showing higher intensities for the iTBS session relative to sham session (EMD = 2.3% MSO [0.0, 4.6], *P* = 0.047). There were no other main effects or interactions for all other baseline stimulation intensities (all *P* > 0.05).

Baseline MEP amplitude for corticospinal excitability and MEP latency differences are shown in Table 2. For PA*_1mV_* MEP amplitude, there was an interaction between session and age (*F*_1,1121_ = 4.194, *P* = 0.041), with *post-hoc* comparisons revealing larger MEP amplitude for young participants relative to older participants (EMD = 0.14 mV [0.02, 0.26], *P* = 0.024). For baseline MEP latency differences, responses differed between coil orientations (*F*_1,112_ = 165.20, *P* < 0.0001), where PA-LM latencies were shorter than AP-LM latencies (EMD = 1.95 ms [1.65, 2.25], *P* < 0.0001), as expected. There were no main effects or interactions for all other baseline MEP amplitude or MEP latency differences (all *P* > 0.05). Absolute MEP amplitude between sessions for young and older adults is presented in Supplementary Materials.

**Table 2.** Baseline responses of corticospinal excitability and I-wave recruitment between sessions.

| Measure | Young |  | Older |  |
| --- | --- | --- | --- | --- |
|  | PMd iTBS-M1 iTBS | PMd sham-M1 iTBS | PMd iTBS-M1 iTBS | PMd sham-M1 iTBS |
| **PA** |  |  |  |  |
| PA-LM latency (ms) | 1.39 [0.89, 1.88] | 1.54 [1.04, 2.03] | 1.97 [1.47, 2.46] | 2.02 [1.52, 2.51] |
| 1mV<sub>PA</sub> (mV) | 1.03 [0.94, 1.12] | 0.93 [0.85, 1.01] | 0.89 [0.81, 0.97]<sup>a</sup> | 0.96 [0.87, 1.04] |
| 0.5mV<sub>PA</sub> (mV) | 0.53 [0.46, 0.60] | 0.49 [0.42, 0.56] | 0.50 [0.43, 0.57] | 0.51 [0.44, 0.58] |
| **AP** |  |  |  |  |
| AP-LM latency (ms) | 3.49 [3.00, 3.98]<sup>b</sup> | 3.54 [3.05, 4.03]<sup>b</sup> | 3.69 [3.20, 4.18]<sup>b</sup> | 3.99 [3.50, 4.48]<sup>b</sup> |
| 1mV<sub>AP</sub> (mV) | 0.97 [0.87, 1.06] | 0.88 [0.79, 0.97] | 1.02 [0.91, 1.14] | 0.99 [0.88, 1.10] |
| 0.5mV<sub>AP</sub> (mV) | 0.47 [0.41, 0.53] | 0.44 [0.39, 0.50] | 0.45 [0.40, 0.51] | 0.45 [0.39, 0.50] |
| Data show EMM [95% CI; lower, upper]. <sup>a</sup>*P* < 0.05 compared to young. <sup>b</sup>*P* < 0.05 compared to PA-LM latency. |  |  |  |  |

### Changes in corticospinal excitability following PMd iTBS

The participants’ perceptions of PMd iTBS and PMd sham are shown in Table 3. While there were no differences between sessions in the extent of discomfort (*t*_29_ = 0.25, *P* = 0.804) or FDI activation (*t*_29_ = 0.10, *P* = 0.918) experienced by the participants, the locality of stimulation differed (*t*_29_ = 3.98, *P* = 0.004), with the sensation of iTBS perceived as more widespread relative to electrical scalp stimulation in sham.

**Table 3.**
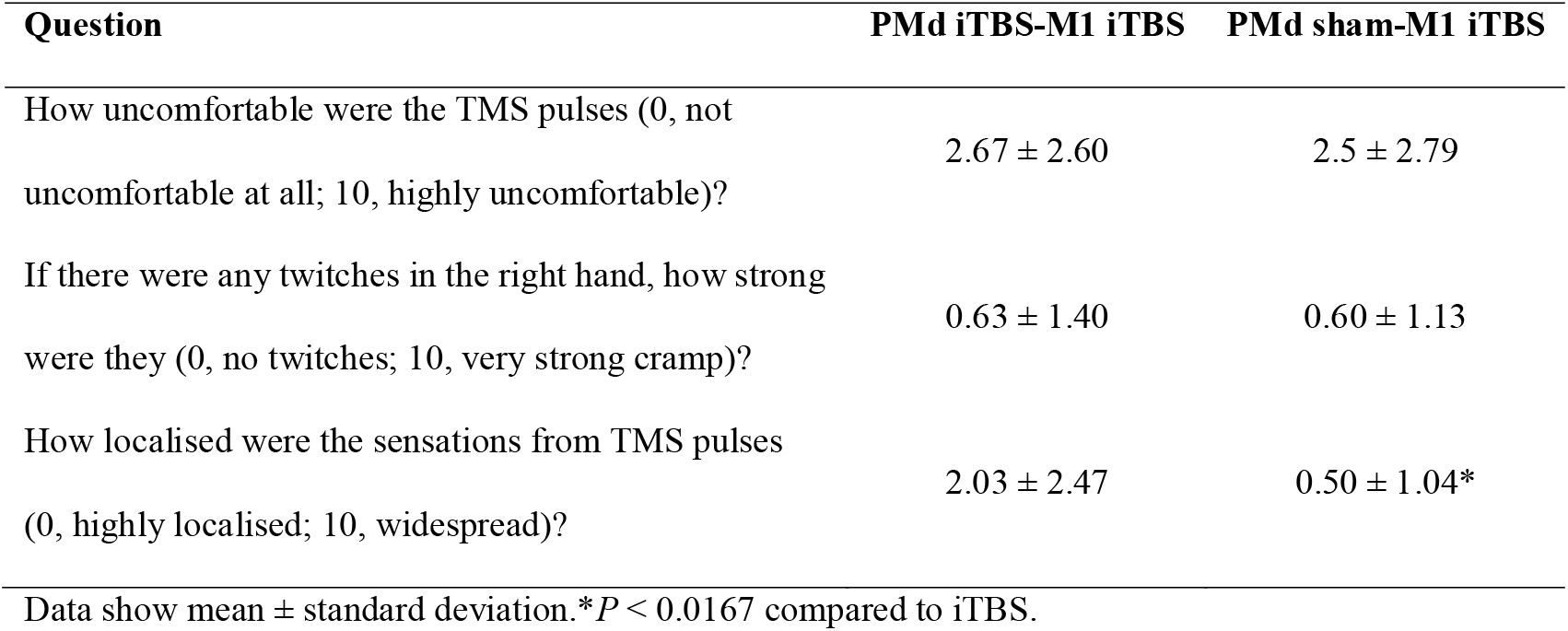
Comparison of VAS responses (mean ± STD) between sessions.

| Question | PMd iTBS-M1 iTBS | PMd sham-M1 iTBS |
| --- | --- | --- |
| How uncomfortable were the TMS pulses (0, not uncomfortable at all; 10, highly uncomfortable)? | $2.67 \pm 2.60$ | $2.5 \pm 2.79$ |
| If there were any twitches in the right hand, how strong were they (0, no twitches; 10, very strong cramp)? | $0.63 \pm 1.40$ | $0.60 \pm 1.13$ |
| How localised were the sensations from TMS pulses (0, highly localised; 10, widespread)? | $2.03 \pm 2.47$ | $0.50 \pm 1.04^*$ |

Data show mean ± standard deviation. \* $P < 0.0167$ compared to iTBS.

Changes in MEP*_1mV_* and MEP*_0.5mV_* measures of corticospinal excitability following PMd iTBS are shown in Figure 2. PA*_1mV_* MEP amplitude did not differ between sessions (*F*_1,1114_ = 0.90, *P* = 0.343; Fig. 2A) or age groups (*F*_1,1114_ = 0.12, *P* = 0.726), and there was no interaction between factors (*F*_1,1114_ = 2.41, *P* = 0.121). AP*_1mV_* MEP amplitude did not vary between sessions (*F*_1,996_ = 2.33, *P* = 0.127; Fig. 2B) or age groups (*F*_1,996_ = 1.31, *P* = 0.252), and there was no interaction between factors (*F*_1,996_ = 0.51, *P* = 0.476). In contrast, while PA*_0.5mV_* MEP amplitude did not differ between age groups (*F*_1,1152_ = 0.11, *P* = 0.740), responses varied between sessions (*F*_1,1152_ = 4.23, *P* = 0.040; Fig. 2C), with increased MEP amplitude following PMd iTBS relative to sham (EMD = 26.3% [0.7, 51.9], *P* = 0.044). There was no interaction between factors (*F*_1,1152_ = 0.11, *P* = 0.741). AP*_0.5mV_* MEP amplitude did not vary between sessions (*F*_1,1073_ = 1.04, *P* = 0.308; Fig. 2D) or age groups (*F*_1,1073_ = 2.80, *P* = 0.095), and there was no interaction between factors (*F*_1,1073_ = 1.03, *P* = 0.310).

**Figure 2.**
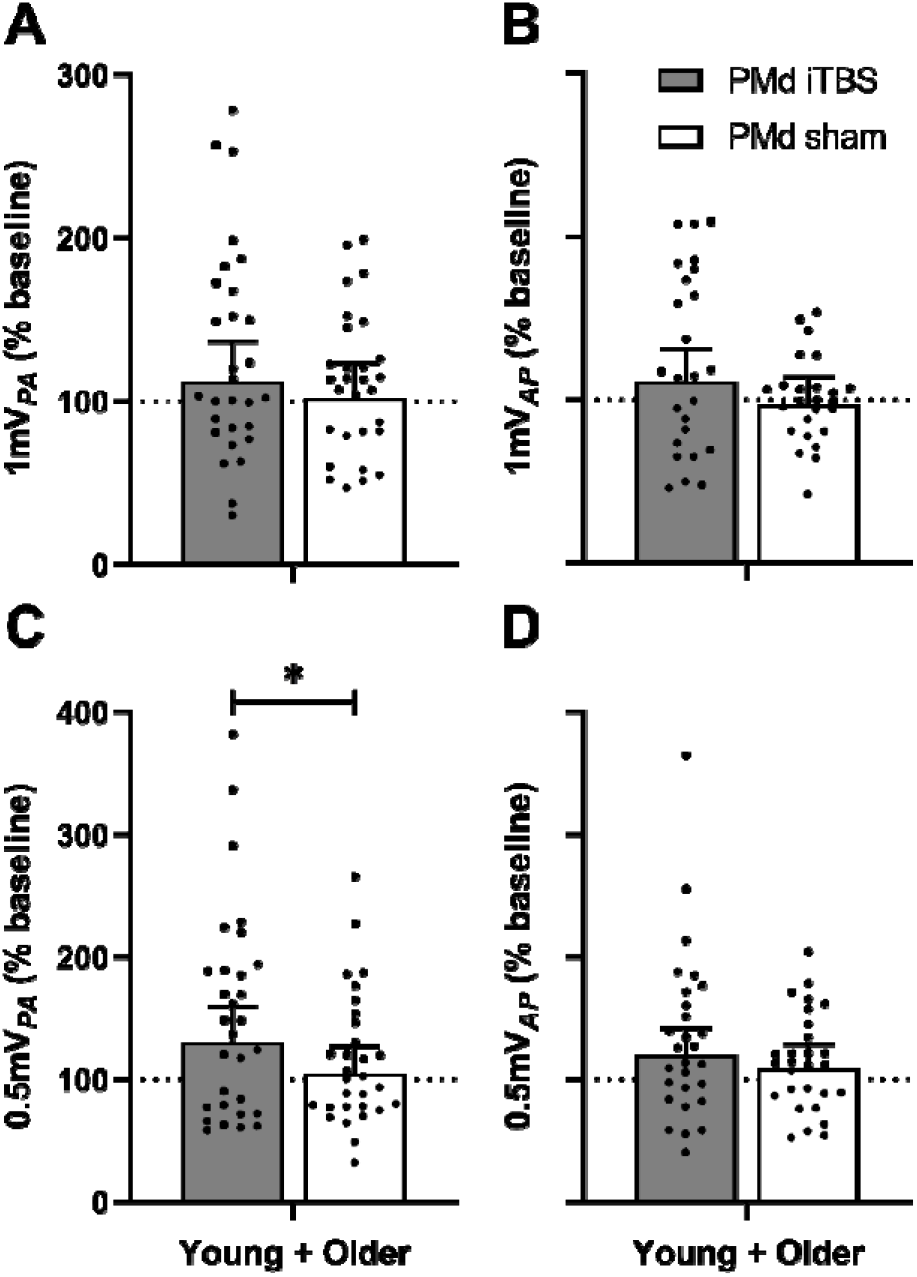
Changes in PA*_1mV_* (A), AP*_1mV_* (B), PA*_0.5mV_* (C), and AP*_0.5mV_* (D) measures of corticospinal excitability following PMd iTBS (grey) and sham (white) stimulation in all participants. Data show EMM (95% CI) with individual subject means. \**P* < 0.05.

### Changes in corticospinal excitability and I-wave recruitment following M1 iTBS

#### Corticospinal excitability

Changes in MEP*_1mV_* measures of corticospinal excitability following PMd iTBS-M1 iTBS and PMd sham-M1 iTBS are presented in Figure 3. PA*_1mV_* MEP amplitude (Fig. 3A) did not vary between sessions (*F*_1,2234_ = 2.20, *P* = 0.138), time points (*F*_1,2234_ = 0.15, *P* = 0.696), or age groups (*F*_1,2234_ = 1.17, *P* = 0.279), and there were no interactions between factors (all *P* > 0.05). AP*_1mV_* MEP amplitude also did not differ between sessions (*F*_1,1921_ = 0.98, *P* = 0.323), time points (*F*_1,1921_ = 1.14, *P* = 0.286), or age groups (*F*_1,1921_ = 1.21, *P* = 0.272), but there was an interaction between session and time (*F*_1,1921_ = 9.94, *P* = 0.002; Fig 3B). *Post-hoc* comparisons showed that MEP amplitude following PMd sham-M1 iTBS was increased at 5 minutes compared to PMd iTBS-M1 iTBS (EMD = 29.7% [6.6, 52.8], *P* = 0.012), and compared to 30 minutes (EMD = 30.3% [7.3, 53.3], *P* = 0.010). There were no other interactions (all *P* > 0.05). In addition, there was no effect of stimulation intensity on MEP amplitude (*F*_1,1921_ = 0.40, *P* = 0.527).

**Figure 3.**
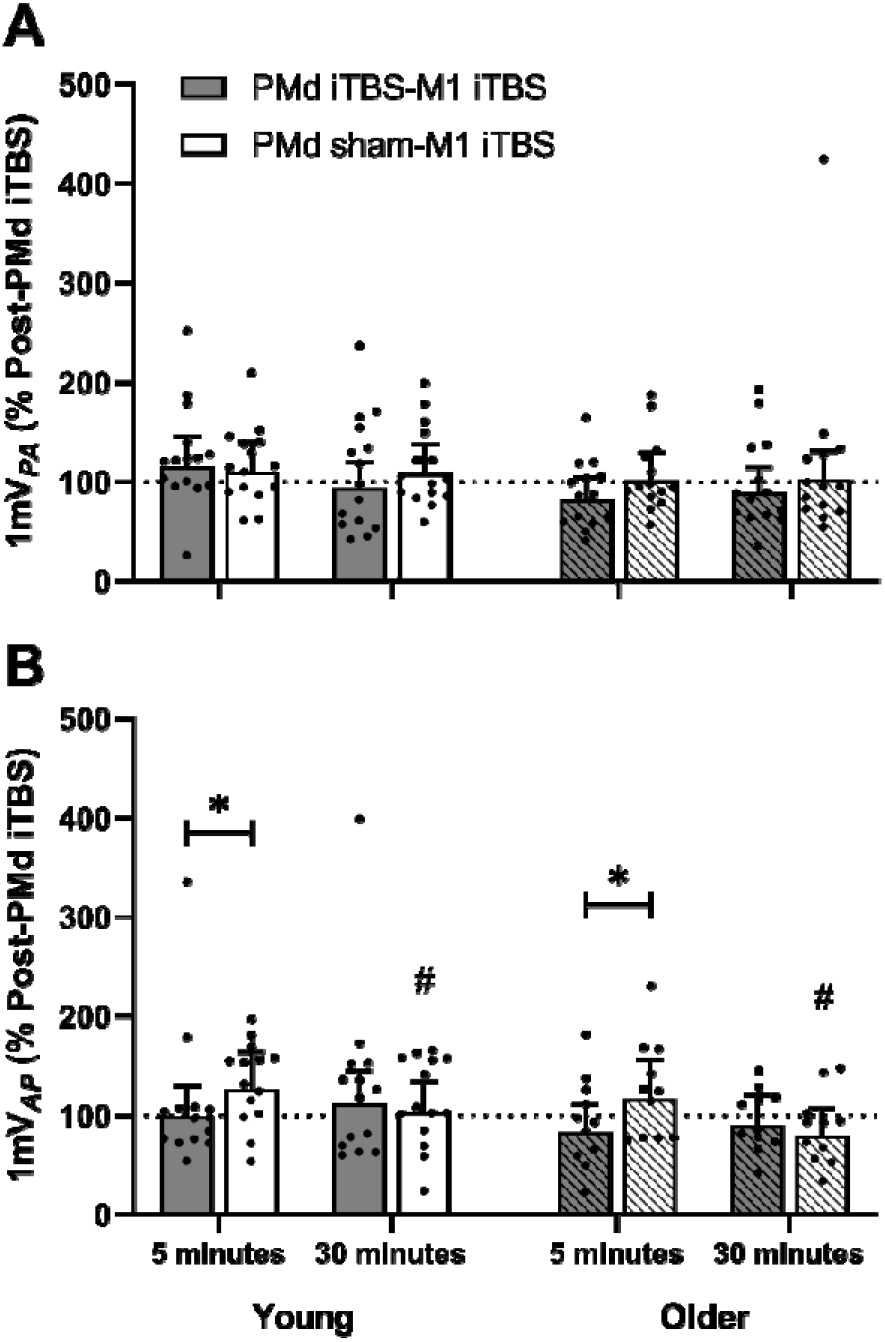
Changes in PA*_1mV_*(A) and AP*_1mV_* (B) measures of corticospinal excitability following PMd iTBS-M1 iTBS (grey) and PMd sham-M1 iTBS (white) in young (no stripes) and older adults (stripes) at 5 and 30 minutes. Data show EMM (95% CI) with individual subject means. \**P* < 0.05. #*P* < 0.05 compared to 5 minutes in same session.

Changes in MEP*_0.5mV_* measures of corticospinal excitability are presented in Figure 4. While PA*_0.5mV_* MEP amplitude did not differ between time points (*F*_1,2311_ = 0.03, *P* = 0.874) or age groups (*F*_1,2311_ = 0.17, *P* = 0.678), responses varied between sessions (*F*_1,2311_ = 17.4, *P* < 0.05), with increased MEP amplitude following PMd sham-M1 iTBS (EMD = 34.3% [17.5, 51.0], *P* < 0.05). Furthermore, there was an interaction between session, time, and age (*F*_1,2311_ = 4.71, *P* = 0.030; Fig. 4A). *Post-hoc* analysis revealed increased MEP amplitude following PMd sham-M1 iTBS compared to PMd iTBS-M1 iTBS for young adults at 30 minutes (EMD = 50.7% [20.1, 81.3], *P* = 0.001), while this effect was observed for older adults at 5 (EMD = 43.0% [13.9, 72.0], *P* = 0.004) and 30 minutes (EMD = 32.0% [3.6, 60.4], *P* = 0.027). For AP*_0.5mV_*, MEP amplitude did not vary between sessions (*F*_1,2141_ = 0.13, *P* = 0.723) or age groups (*F*_1,2141_ = 3.12, *P* = 0.077) (Fig. 4B). However, responses differed between time points (*F*_1,2141_ = 5.91, *P* = 0.015; Fig. 4C), with *post-hoc* analysis revealing that MEP amplitude was increased at 5 minutes relative to 30 minutes post-M1 iTBS (EMD = 22.0% [3.9, 40.0], *P* = 0.017). There were no interactions between factors (all *P* > 0.05).

**Figure 4.**
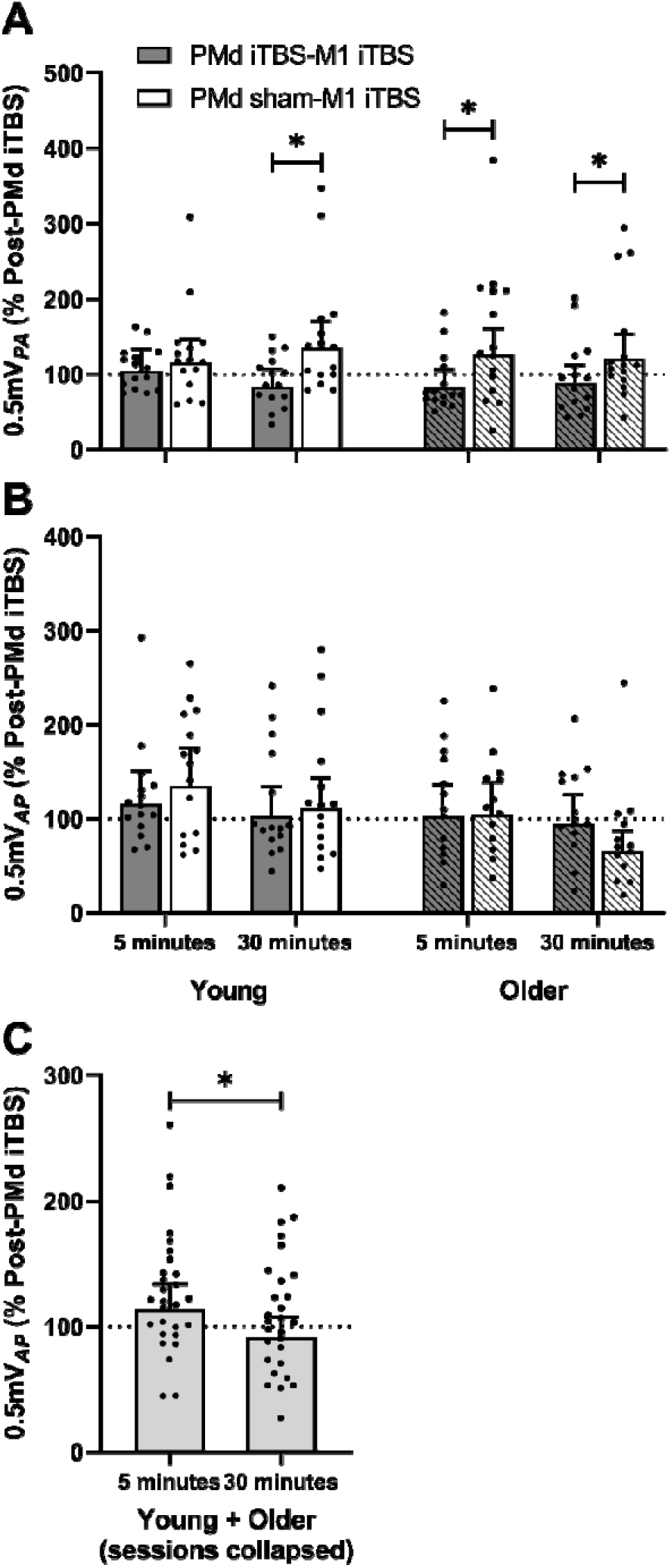
Changes in PA*_0.5mV_*(A) and AP*_0.5mV_* (B) measures of corticospinal excitability following PMd iTBS-M1 iTBS (grey) and PMd sham-M1 iTBS (white) in young (no stripes) and older adults (stripes) at 5 and 30 minutes. (C) Changes in AP*_0.5mV_* following M1 iTBS in all participants (light grey) at 5 and 30 minutes. Data show EMM (95% CI) with individual subject means. \**P* < 0.05.

#### I-wave recruitment

There was no difference between sessions (*F*_1,112_ = 0.72, *P* = 0.399), coil orientations (*F*_1,112_ = 0.09, *P* = 0.766), or age groups (*F*_1,112_ = 0.38, *P* = 0.538), and there were no interactions between factors (all *P* > 0.05).

#### Correlation analyses

Baseline PA-LM and AP-LM latencies were not related to changes in single-pulse measures of corticospinal excitability (PA*_1mV_*, AP*_1mV_*, PA*_0.5mV_*, AP*_0.5mV_*) following PMd iTBS (all *P* > 0.05). Baseline PA-LM and AP-LM latencies were not related to changes in single-pulse measures of corticospinal excitability following PMd sham-M1 iTBS (all *P* > 0.05). In contrast, while changes in AP*_1mV_* MEP amplitude following PMd iTBS were not related to changes in AP*_1mV_* responses following M1 iTBS (ρ = −0.361, *P* = 0.076; Fig. 5B), changes in PA*_1mV_*, PA*_0.5mV_*, and AP*_0.5mV_* MEP amplitude following PMd iTBS were negatively correlated with changes in PA*_1mV_* (ρ = 0.-577, *P* = 0.001; Fig. 5A), PA*_0.5mV_* (ρ = −0.616, *P* = 0.0003; Fig. 5C), and AP*_0.5mV_* (ρ = −0.551, *P* = 0.002; Fig. 5D) responses following M1 iTBS, respectively.

**Figure 5.**
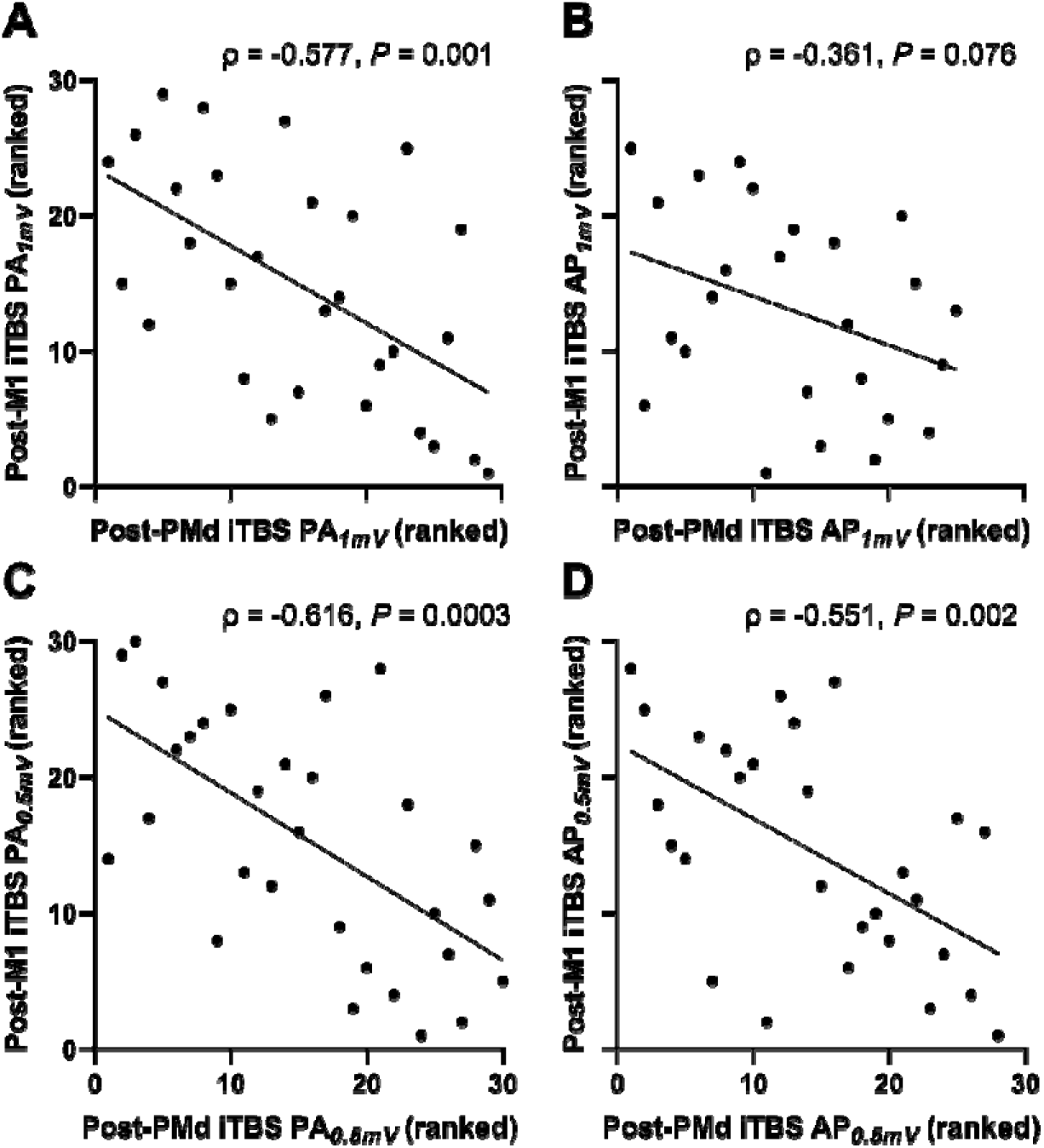
Correlation of ranked changes in post-PMd iTBS measures of corticospinal excitability (PA*_1mV_*, A; AP*_1mV_*, B; PA*_0.5mV_*, C; AP*_0.5mV_*, D) with ranked changes in post-M1 iTBS measures of corticospinal excitability.

## Discussion

In the present study, we investigated the influence of PMd on the plasticity of early and late I-wave-generating circuits in M1 of young and older adults. This was achieved by applying PMd iTBS as a priming intervention to modify the neuroplastic response of M1 to subsequent iTBS (PMd iTBS-M1 iTBS, PMd sham-M1 iTBS). We measured changes in corticospinal excitability (PA*_1mV_*, AP*_1mV_*, PA*_0.5mV_*, AP*_0.5mV_*) and I-wave recruitment (PA-LM latency, AP-LM latency) following the intervention. The findings show that PMd iTBS specifically modulated the excitability of the early I-wave circuits in both young and older adults.

Moreover, PMd iTBS disrupted the neuroplastic response of the early I-wave circuits to M1 iTBS in both young and older adults, whereas the neuroplastic response of the late I-wave circuits was unaffected in both age groups.

### PMd influence on corticospinal excitability in young and older adults

Previous work has reported that application of iTBS to PMd facilitates PA*_1mV_* measures of M1 corticospinal excitability in young adults by ∼30%, which is thought to stem from the induction of LTP-like effects within PMd, resulting in increased excitability within M1 (Meng *et al*., 2020). Furthermore, we have demonstrated previously that this effect on PA*_1mV_* is preserved with ageing and extends to AP*_1mV_* measures of corticospinal excitability (Liao *et al*., 2023). The absence of any changes in PA*_1mV_*or AP*_1mV_* within the present study is therefore inconsistent with these previous findings. However, inter- and intraindividual variability in the changes in M1 excitability following TBS is well-documented (Hamada *et al*., 2013; Corp *et al*., 2020; Guerra *et al*., 2020). In particular, there is some variability in the time course of facilitation following PMd iTBS. For example, one study reported that the facilitation of MEP amplitude only occurred at 15 minutes (Meng *et al*., 2020), whereas we previously demonstrated facilitation of MEP amplitude that persisted from 5 to 40 minutes following PMd iTBS (Liao *et al*., 2023). Consequently, our decision to record MEPs at 5 minutes post-PMd iTBS may have limited the ability to detect changes in corticospinal excitability due to the priming intervention.

Although PA*_1mV_* and AP*_1mV_* MEP amplitude were not modulated following PMd iTBS, PA*_0.5mV_* was facilitated (by ∼30%) for both young and older adults. The conventional interpretation of how TMS intensity and current direction influence I-wave recruitment suggests that low-intensity PA TMS preferentially recruits early I-waves, whereas low-intensity AP TMS preferentially recruits late I-waves (Hamada *et al*., 2013), with either current direction able to recruit both I-waves as the stimulation intensity is increased (Di Lazzaro *et al*., 2001; Di Lazzaro *et al*., 2003). We therefore applied single-pulse TMS at relatively lower intensities compared to MEP*_1mV_* (PA*_0.5mV_*, AP*_0.5mV_*), where PA*_0.5mV_* is likely more selective for activation of the early I-waves, while AP*_0.5mV_* is likely more selective for the late I-waves (Opie *et al*., 2022). Given that we previously reported potentiation of both PA*_0.5mV_* and AP*_0.5mV_* (by ∼50-100%) following PMd iTBS (Liao *et al*., 2023), the increase in PA*_0.5mV_* within the present study suggests that the effect of PMd iTBS on early I-wave excitability may be immediate and more consistent. Importantly, previous work has shown that PMd iTBS applied as it was in the current study is unlikely to have activated M1 directly. Specifically, Huang and colleagues (2009) assessed the intensity required to activate M1 when TMS was applied over PMd, and showed that 80% of this (matching the level applied during iTBS) applied to M1 does not influence M1 excitability (Huang *et al*., 2009). Given that we located PMd using similar methods (Huang *et al*., 2009; Huang *et al*., 2018; Meng *et al*., 2020), it is therefore unlikely that PMd iTBS activated M1 directly in the present study.

Despite the present findings demonstrating that PMd iTBS increased early I-wave excitability, this effect was not different between young and older adults, suggesting that the influence of PMd on early I-wave excitability may be preserved with ageing. This contrasts with our previous work, which specifically demonstrated weakened direct PMd modulation of early I-waves in older adults (Liao *et al*., 2023). Given that both studies employed the same methods to assess changes in M1 excitability following PMd iTBS, participant factors such as genetics, pharmacology, aerobic exercise, and diet that are known to influence cortical plasticity (Ridding & Ziemann, 2010; Phillips, 2017) may have confounded the present findings. As the contributions of participant characteristics on PMd-M1 communication were not examined in the present study, and the small sample sizes were not powered for such subanalyses, it will be important to characterise their involvement in future studies.

### PMd influence on M1 plasticity in young and older adults

Previous work in young participants demonstrated that applying continuous TBS (cTBS) to PMd disrupts the neuroplastic response of M1 to both iTBS and cTBS, assessed using PA*_1mV_* MEPs (Huang *et al*., 2018). This demonstrated that LTP- and LTD-like effects within M1 can be modulated by PMd cTBS, which was thought to arise from heterosynaptic metaplastic effects, where the modulation of local synaptic plasticity within PMd affected subsequent changes in remote synapses (that were not initially activated) within M1 (Huang *et al*., 2018). In the present study, we demonstrated that applying iTBS to PMd also disrupts the LTP-like effects of M1 iTBS for AP*_1mV_* measures of corticospinal excitability. However, given that iTBS produces LTP-like effects while cTBS produces LTD-like effects, this disruption of AP*_1mV_* facilitation may stem from a different mechanism more consistent with homeostatic metaplasticity (Müller *et al*., 2007; Todd *et al*., 2009; Murakami *et al*., 2012). Importantly, this response did not differ between young and older adults, suggesting that the influence of PMd on the plasticity of AP circuits within M1 is maintained with age. In addition, this difference is also unlikely to be driven by AP*_1mV_* stimulation intensity differences at baseline, as investigation of this confounding factor did not reveal any effects on post-intervention AP*_1mV_* MEP amplitude.

Furthermore, PMd iTBS disrupted the effects of M1 iTBS on PA*_0.5mV_* (early), but not AP*_0.5mV_* (late) circuits. This suggests that the influence of PMd on M1 plasticity is specific to the early I-waves, which is unexpected given that previous work has demonstrated a stronger influence of PMd on late I-wave circuits (Volz *et al*., 2015; Aberra *et al*., 2020; Liao *et al*., 2023). One possible explanation is that shorter AP MEP onset latencies (more consistent with early I-wave recruitment) have been reported to predict stronger premotor-M1 functional connectivity (Volz *et al*., 2015) and the nature of this communication may contribute to the influence of PMd on M1 plasticity. This appears consistent with our AP*_1mV_* findings, which may have occurred as the higher stimulus intensity required to record AP*_1mV_* resulted in mixed recruitment of early and late I-waves, but that changes in AP*_1mV_* were driven specifically by the early I-waves (Di Lazzaro *et al*., 2001; Di Lazzaro *et al*., 2003; Liao *et al*., 2022). This is further complemented by the correlation analysis results demonstrating that larger facilitation of PA*_0.5mV_* post-PMd iTBS is correlated with smaller facilitation of PA*_0.5mV_*post-M1 iTBS, suggesting that this homeostatic metaplastic effect is likely related to the early I-wave circuits. While a similar correlation was also shown for PA*_1mV_* and AP*_0.5mV_*, PMd iTBS-M1 iTBS did not disrupt the potentiation of these measures when compared to PMd sham-M1 iTBS session. It is possible that the higher stimulus intensities required for PA*_1mV_* and AP*_0.5mV_* (relative to PA*_0.5mV_*) may have also resulted in mixed recruitment of early and late I-waves (Liao *et al*., 2022). In particular, given that there is growing evidence to suggest that PA and AP TMS can activate distinct populations of early and late I-waves (i.e., PA- and AP-sensitive early and late I-waves) (Spampinato *et al*., 2020; Opie & Semmler, 2021), PA*_1mV_* and AP*_0.5mV_* may have recruited other I-wave circuits that were less sensitive to the modulatory effects of iTBS. However, this will need to be clarified in future research using techniques that are more selective to these different I-waves, such as modifying the TMS pulse width (Hannah & Rothwell, 2017). Despite this, we provide new evidence that PMd iTBS specifically modulates M1 plasticity of early I-wave circuits recruited by AP stimulation.

While M1 iTBS in isolation (PMd sham-M1 iTBS) potentiated PA*_0.5mV_* responses (compared with PMd iTBS-M1 iTBS) in both age groups, the timing of this response varied between groups. Whereas differences between sham and real PMd iTBS sessions were immediate for older adults, they were only apparent after 30 minutes in young adults. Given that M1 iTBS has not been shown to differentially modulate corticospinal excitability in young and older adults (Di Lazzaro *et al*., 2008; Young-Bernier *et al*., 2014; Dickins *et al*., 2015; Opie *et al*., 2017), this outcome seems unlikely to reflect effects of age within M1. An alternative explanation could be that the modulatory effects of PMd iTBS differed between groups, with younger adults having a stronger response that was more resistant to the subsequent effects of M1 iTBS. This is supported by the amplitude of PA*_0.5mV_* being reduced 5 minutes after M1 iTBS in older, but not young adults in the session involving real PMd iTBS (Fig. 4A).

Although speculative, this outcome would be consistent with our previous finding that the influence of PMd iTBS on PA*_0.5mV_* is reduced in older adults (Liao *et al*., 2023). However, this speculation will require additional studies that more effectively characterise the time course of facilitation in young and older adults. For example, previous work investigating the effects of PMd cTBS on M1 neuroplastic response to iTBS or cTBS monitored changes in corticospinal excitability for two hours following PMd cTBS (during which excitability returned to baseline levels) before applying subsequent M1 iTBS or cTBS (Huang *et al*., 2018).

### PMd and M1 influence on I-wave recruitment in young and older adults

The ability to recruit both early and late I-waves can be investigated by comparing the latencies evoked by PA and AP TMS to the latencies of direct corticospinal activation (PA-LM, early; AP-LM, late) (Hamada *et al*., 2013). The prototypical values for these measures reveal shorter PA-LM latencies (∼1.5 ms) compared to AP-LM latencies (∼3 ms), providing an index of early and late I-wave recruitment, respectively (Hamada *et al*., 2013).

Importantly, previous studies have shown that the ability to recruit late I-waves with AP TMS predicts the neuroplastic response of M1 to iTBS (Hamada *et al*., 2013; Volz *et al*., 2019), with AP inputs thought to originate from PMd (Volz *et al*., 2015; Aberra *et al*., 2020). It has also been demonstrated that AP-LM latencies can be shortened using M1 iTBS, which was suggested to reflect the direct modulation of the late I-wave circuitry (Volz *et al*., 2019).

Although we also assessed changes in PA-LM and AP-LM latencies following PMd sham-M1 iTBS in the present study, the intervention failed to modulate the I-wave latencies. It is possible that changes in AP-LM latencies occur immediately following iTBS, as the MEP latency measures were recorded at least 45 minutes either side of PMd and M1 iTBS in order to avoid complications involving the effects of muscle activation on neuroplasticity responses (Huang *et al*., 2008; Thirugnanasambandam *et al*., 2011; Goldsworthy *et al*., 2015). Consequently, the effects of M1 iTBS on I-wave latencies will have to be clarified in future studies.

Importantly, baseline I-wave recruitment was not correlated with changes in corticospinal excitability following M1 iTBS in isolation, in contrast to previous findings (Hamada *et al*., 2013; Volz *et al*., 2019). While the difference between the present study and previous studies is that we included older participants, no differences between age groups were shown for I-wave recruitment or corticospinal excitability in the present study. The variability in the present findings may therefore involve contributions from other factors. For example, recent work assessing variability of M1 iTBS has suggested that the ability of iTBS to engage neural oscillations in the β range (13-30 Hz) may be an important predictor of the neuroplastic response to iTBS (Leodori *et al*., 2021). Enhancing premotor-M1 communication using cortico-cortical paired associated stimulation (ccPAS) has been recently shown to improve the synchronisation of neural oscillations (which is thought to mediate neuronal communication and plasticity) in the β range (Trajkovic *et al*., 2023). Further investigation involving these measures may therefore better characterise the variability of iTBS, and may also have applications in understanding PMd-M1 communication.

In conclusion, the application of iTBS over PMd potentiated corticospinal excitability and disrupted the effects of subsequent M1 iTBS. Specifically, our results show that PMd may more consistently influence the excitability of early I-waves in young and older adults. Importantly, we provide new evidence that PMd disrupts M1 plasticity of early I-wave circuits in both age groups. It will therefore be useful in future studies to investigate how PMd modulation of M1 plasticity influences different feature of motor skill learning in young and older adults.

