## Supplemental Table 1. for "Modulation of dorsal premotor cortex disrupts neuroplasticity of primary motor cortex in young and older adults"

**Supplementary Material**

**Results**

| **Table 1. MEP amplitude between TMS sessions for young and older adults at baseline, Post-PMd-iTBS, 5 minutes post-M1-iTBS, and 30 minutes post-M1 iTBS** | | | | | | |
| --- | --- | --- | --- | --- | --- | --- |
| **Measure** | **Age** | **Session** | **Baseline** | **Post-PMd iTBS** | **5 minutes post-M1 iTBS** | **30 minutes post-M1 iTBS** |
| **PA** |  |  |  |  |  |  |
| 1mV*_PA_* (mV) | Young | PMd iTBS-M1 iTBS | 1.03 ± 0.77 | 1.13 ± 0.87 | 1.27 ± 0.97 | 1.08 ± 0.86 |
|  |  | PMd sham-M1 iTBS | 0.90 ± 0.53 | 1.18 ± 0.86 | 0.99 ± 0.76 | 1.04 ± 0.70 |
|  | Older | PMd iTBS-M1 iTBS | 0.93 ± 0.63 | 1.05 ± 0.89 | 1.12 ± 0.72 | 1.14 ± 0.79 |
|  |  | PMd sham-M1 iTBS | 0.97 ± 0.75 | 0.96 ± 0.76 | 1.03 ± 1.03 | 1.00 ± 0.77 |
| 0.5mV*_PA_* (mV) | Young | PMd iTBS-M1 iTBS | 0.54 ± 0.48 | 0.77 ± 0.84 | 0.79 ± 0.72 | 0.66 ± 0.62 |
|  |  | PMd sham-M1 iTBS | 0.52 ± 0.45 | 0.78 ± 0.73 | 0.62 ± 0.53 | 0.63 ± 0.60 |
|  | Older | PMd iTBS-M1 iTBS | 0.50 ± 0.47 | 0.59 ± 0.66 | 0.63 ± 0.58 | 0.75 ± 0.74 |
|  |  | PMd sham-M1 iTBS | 0.53 ± 0.50 | 0.58 ± 0.64 | 0.79 ± 0.80 | 0.70 ± 0.60 |
| **AP** |  |  |  |  |  |  |
| 1mV*_AP_* (mV) | Young | PMd iTBS-M1 iTBS | 0.97 ± 0.79 | 1.04 ± 0.85 | 1.14 ± 1.17 | 1.20 ± 1.04 |
|  |  | PMd sham-M1 iTBS | 1.02 ± 0.72 | 1.38 ± 0.97 | 1.13 ± 0.90 | 1.18 ± 0.80 |
|  | Older | PMd iTBS-M1 iTBS | 0.88 ± 0.79 | 0.84 ± 0.71 | 1.08 ± 0.94 | 0.97 ± 1.00 |
|  |  | PMd sham-M1 iTBS | 0.99 ± 0.87 | 1.02 ± 0.89 | 1.32 ± 1.27 | 0.88 ± 0.84 |
| 0.5mV*_AP_* (mV) | Young | PMd iTBS-M1 iTBS | 0.53 ± 0.54 | 0.66 ± 0.75 | 0.74 ± 0.84 | 0.74 ± 0.97 |
|  |  | PMd sham-M1 iTBS | 0.52 ± 0.42 | 0.71 ± 0.72 | 0.73 ± 0.57 | 0.67 ± 0.58 |
|  | Older | PMd iTBS-M1 iTBS | 0.54 ± 0.51 | 0.52 ± 0.62 | 0.76 ± 0.79 | 0.60 ± 0.73 |
|  |  | PMd sham-M1 iTBS | 0.53 ± 0.54 | 0.69 ± 0.73 | 0.76 ± 0.81 | 0.53 ± 0.54 |

Data are presented as mean ± standard deviation.
